# The interactome of histone deacetylase HDA19 in dark-grown Arabidopsis seedlings

**DOI:** 10.1101/2023.09.19.558523

**Authors:** Isabel Cristina Vélez-Bermúdez, Wolfgang Schmidt

## Abstract

Here, we describe a data set derived from an immunoprecipitation (IP)-based analysis of the histone deacetylase HDA19 interactome in etiolated Arabidopsis seedlings. HISTONE DEACETYLASE 19 (HDA19) belongs to the class 1 of the Reduced Potassium Dependence3/Histone Deacetylase-1 (RPD3/HDA1) superfamily and is possibly the most intensively studied HDA. Here, we describe a data set derived from an immunoprecipitation (IP)-based analysis of the histone deacetylase HDA19 interactome in etiolated Arabidopsis seedlings. We believe that this data set presented here provides a valuable resource for follow-up research on novel interacting partners of this central protein.

## Introduction

As a central epigenetic modification, histone acetylation affects the expression of genes with a wide range of functions across all life forms (Shen et al., 2015). Histone acetylation is mediated by histone acetylases and generally promotes DNA-templated transcriptional activity. Histone deacetylases (HDAs) reverse this process, leading to an inactive chromatin state and generally decreased transcriptional activity. Acetylation and deacetylation are central regulatory switches that govern responses to various environmental cues, orchestrating chromatin dynamics and gene activity to modulate the phenotypic readout.

The Arabidopsis genome harbours 18 HDAs, which are organized in three superfamilies comprised of the HDA classes 1-4 (Seto and Yoshida, 2014). HISTONE DEACETYLASE 19 (HDA19) belongs to the class 1 of the Reduced Potassium Dependence3/Histone Deacetylase-1 (RPD3/HDA1) superfamily and is possibly the most intensively studied HDA. HDA19 has reported roles across a broad landscape of processes and was shown to be a crucial player in seed development (Long et al., 2006; Zhou et al., 2020; Chen et al., 2023), pathogen response (Zhou et al., 2005), phosphate deficiency (Chen et al., 2015), light signalling (Jing et al., 2020), the regulation of flowering time (Ning et al., 2019), hormone responses (Kim, 2019; Mehdi et al. 2016), and floral organ identity genes (Krogan et al., 2012; Bollier et al., 2018; Ning et al., 2019). Similar to other RPD3-type HDACs, HDA19 forms various complexes that play pivotal roles in various stress responses (Feng et al., 2021; Vélez-Bermúdez and Schmidt, 2021), some of which possess yet unexplored stress-specific compositions.

In this Data Report, we provide an immunoprecipitation (IP)-based analysis of the HDA19 interactome in etiolated Arabidopsis seedlings. We believe that the interactome presented here provides a valuable resource for follow-up research on novel interacting partners of this central protein. The material used for this experiment was derived from hypocotyls of 6-day-old etiolated Arabidopsis seedlings. The dataset contains a total of 6 files, 3 independent biological replicates of each Col-0 (control plants) and 35S::HDA19-GFP plants.

## Materials and methods

### Plant materials and growth conditions

*Arabidopsis thaliana* Col-0 and the transgenic line 35S::HDA19-GFP were used in this study. 35S::HDA19-GFP lines have been described previously (Zhou et al., 2005). Seeds were soaked in 35% bleach for 5□ min, washed five times for 5 min with sterile water, and resuspended in 1 mL of sterile water for further use. Seeds were subsequently placed on a growth medium (Estelle and Somerville, 1987; ES medium) containing 5□mM KNO_3_, 2□mM MgSO_4_, 2□mM Ca (NO_3_)_2_, 2.5□mM KH_2_PO_4_, 70□μM H_3_BO_3_, 14□μM MnCl_2_, 1□μM ZnSO_4_, 0.5□μM CuSO_4_, 0.01□μM CoCl_2_, 0.2□μM Na_2_MoO_4_, and 40□μM Fe-EDTA, solidified with 0.4% Gelrite Pure. MES (1 g/L) and 1.5% (w/v) sucrose were added and the pH was adjusted to 5.5 with KOH. Seeds were stratified on plates for 2 days at 4°C in the dark and grown at 22°C in vertical position in the dark with 70% relative humidity.

### Immunoprecipitation

One gram of hypocotyls from 5-day-old etiolated seedling was used to perform the IP. Experiments were carried out with the μMACS Epitope GFP tag protein isolation kit (MACSmolecular) following the manufacturer’s instructions with minor modifications. The hypocotyls were grounded with liquid nitrogen and resuspended in 500 μL of extraction buffer (50 mM Tris/HCl, pH 7.5, 150 mM NaCl, 1% Triton X-100, 2X complete protease inhibitor cocktail EDTA-free (ROCHE), 1 mM PMSF, and 50 μM MG132). The samples were incubated on ice for 30 minutes with occasional mixing and centrifuged for 20 minutes at 10,000 x g at 4°C. The supernatants were individually collected in fresh tubes, and 400 μL of each input was added to 50 μL of anti-GFP microbeads and incubated for 1 hour and 30 minutes in a mixer set to 60 rpm at 4°C, while 100 μL of each input was kept to be used for Western blots using anti-GFP as a control. The samples were eluted in 50 μL denaturing elution buffer supplied with the kit.

### On-bead trypsin digestion and protein identification

Eluted IP samples were resuspended in 30 µL elution buffer (5% SDS in 50 mM TEAB, pH 8.5), 1.6 μL of 200 mM TCEP and 1.6 μL of 800 mM CAA were added, and the samples were incubated at 45 for 15 min. After alkylation, 3.3 µL of 55.5% phosphoric acid (PA) was added and the pH (~1) was controlled by means of pH paper. After acidification, each sample was mixed with 198 μL of binding buffer (100 mM TEAB in 90% MeOH). The samples were then placed into an S-trap micro in a 1.7 μL receiver tube for waste flow-through, loaded onto an S-trap column and centrifuged at 4,000 g for 2 min to trap the proteins. The samples were then washed in the column with 150 µL of binding buffer and centrifuged three times at 4,000 g for 2 min. An additional centrifugation step (4,000 g for 2 min) was added to fully remove residual binding buffer. The S-trap column was transferred to a fresh 1.7 μL sample tube for the digestion, and 20 µL of protease solution (Lys-C + trypsin, 50 mM TEAB) was added into individual S-traps containing the samples. The cap of each S-trap was loosely closed to limit evaporative loss, and the samples were incubated for 2.5 h at 47 . The columns were removed from the incubator, 40 µL of 50 mM TEAB was added to each column, and the columns were centrifuged at 4,000 g for 1 min. A total of 40 µL of elution buffer 2 (0.2% formic acid in H_2_O) was added onto the column, and the column was centrifuged at 4000 g for 2 min. After this step, 40 µL of elution buffer 3 (50% ACN in H2O) was added onto the column and centrifuged at 4,000 g for 2 min. The samples were dried by speed vacuum and desalted in a ZipTip pipette tip. The dried pellet was then re-dissolved in 10 µL of 0.1% formic acid for LC-MS/MS analysis.

Liquid chromatography was performed by injecting 3 µg of sample in the LC-nESI-Q Exactive mass spectrometer model (Thermo Fisher Scientific) coupled with an on-line nanoUHPLC (Dionex UltiMate 3000 Binary RSLCnano). The Acclaim PepMap 100 C18 trap column (75 µm x 2.0 cm, 3 µm, 100 Å, Thermo Scientific) and the Acclaim PepMap RSLC C18 nano LC column (75 µm x 25 cm, 2 µm, 100 Å) were used to deliver solvent and separate tryptic peptides with a linear gradient from 5% to 35% of acetonitrile in 0.1% (v/v) formic acid for 60 min at a flow rate of 300 nl/min. The acquisition cycle for MS data was performed in the data-dependent mode with a full survey MS scan followed by 10 MS/MS scans of the top 10 precursor ions from the scan. The mass spectrometer was operated in full scan mode (*m*/*z* 350-1,600) in the Orbitrap analyser at a resolution of 70,000. The data dependent MS/MS acquisitions were performed with a 2 *m*/*z* isolation window, 27% NCE (normalized collision energy), and 17,500 resolving power as described previously (Chen et al., 2023).

### Data analysis and identification of putative interactors

Raw data were analysed with the Proteome Discoverer™ Software 2.2 (Thermo Fisher) using the Sequest search algorism. The Arabidopsis protein database (Araport11) was used to conduct the searches; only high confidence proteins were used for the analysis. All peptide spectrum matches were filtered with a q-value threshold of 0.05 (5% FDR), and the proteins were filtered with high confidence threshold (0.05 q-value, 5% FDR). Nuclear proteins identified in more of two biological replicates were considered as putative interacting partners of HDA19.

### Gene ontology

Gene Ontology (GO) analysis was conducted using the ArgiGO v2.0 toolkit web-server (Tian et al., 2017). Significantly enriched GO categories were visualised using REVIGO (Supek et al., 2011) as previously described (Vélez-Bermúdez and Schmidt, 2021).

### Protein-protein interaction (PPI) network

The PPI network was constructed using STRING (https://string-db.org). Only the closest partners of HDA19 proteins were considered.

## Results

The identification of HDA19 protein partners via IP-nano-HPLC-MS/MS relies on the capacity to distinguish true interactors from non-specific binders. To produce the current IP dataset, we used a powerful system to reduce the background in the IP samples. Samples from *Arabidopsis thaliana* Col-0 wild-type and 35S::HDA19_GFP plants were immunoprecipitated using the MultiMACS GFP isolation kit system with µMACS MicroBeads and conjugated to an anti-GFP monoclonal antibody for faster and effective magnetic labelling of GFP-tagged fusion proteins. The complete procedure is depicted in Figure 1A. The IP samples were digested, subjected to nano-HPLC-MS/MS analysis, and HDA19-binding proteins were identified using the Proteome Discoverer software. The dataset provided a total of 371 putative interactors that were identified with high confidence (Supplementary Data File 1). Considering only candidate proteins that are preferentially or exclusively located to the nucleus according to experimental results or predictions, a subset of 52 putatively interacting proteins was identified in at least in 2 biological replicates (Table 1). A GO enrichment analysis of this subset revealed that besides predicted processes such as ‘histone modification’, ‘chromatin organization’, and ‘negative regulation of transcription’, proteins in the categories ‘multicellular organism development’, ‘cell cycle’, ‘protein modification’ and ‘reproduction’ were overrepresented (Figure 2B).

**Figure 1.**
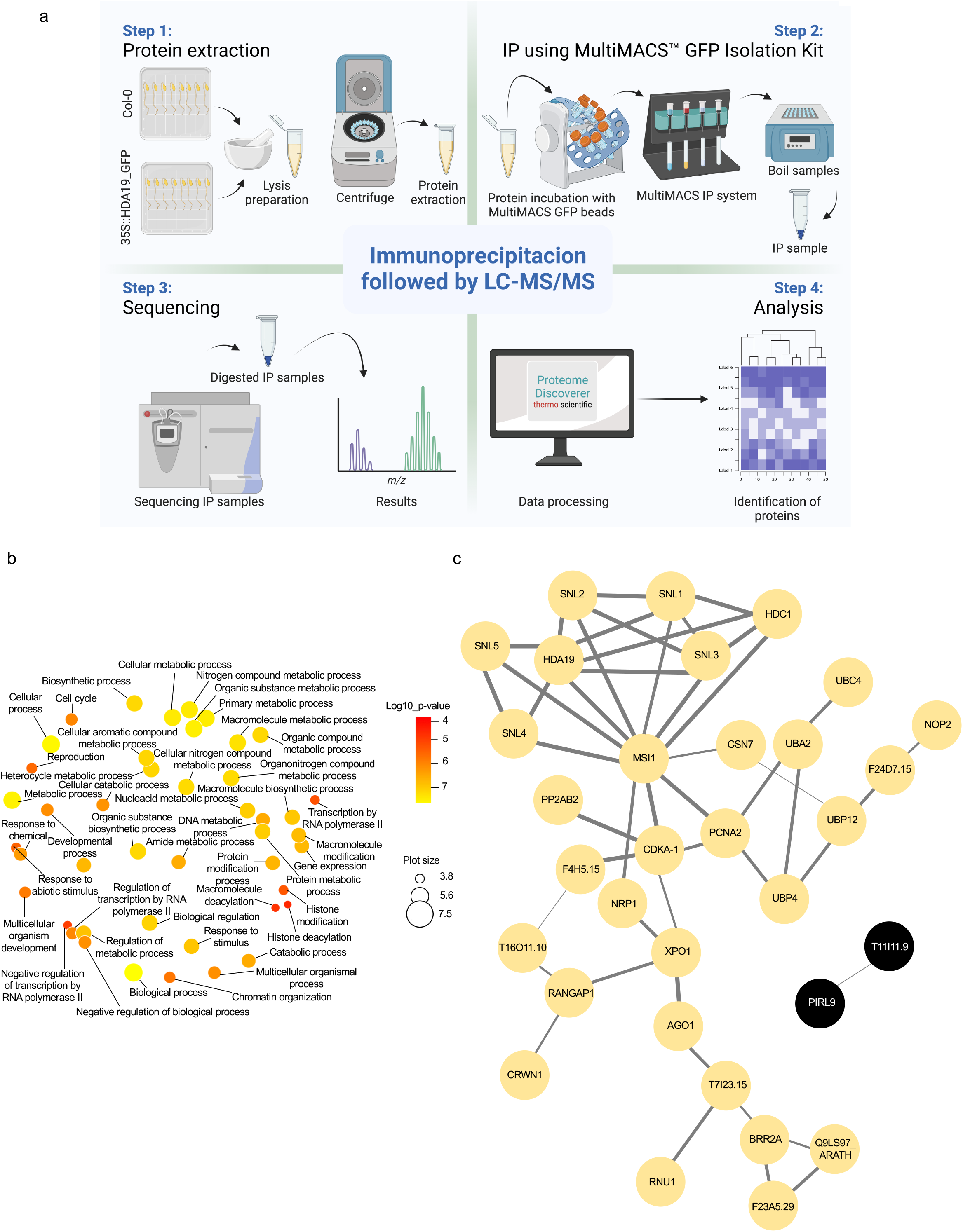
Identification of HDA19-interacting proteins. (A) Experimental flow of the immunoprecipitation analysis. (B) Overrepresented GO categories of putative nuclear-localised HDA19 interactors. Plot color indicates the log10 *P* value of enrichment, the size indicates the frequency of the GO term in the underlying GO annotation database (plots of more general terms are larger). (C) PPI network of putative nuclear-localised HDA19 interactors.

**Table 1.** Putative nuclear-localized binding partners of HDA19. Localization of the proteins was gathered from published experimental evidence or prediction inferred from the Subcellular Location of Proteins in Arabidopsis Database (SUBA).

| TAIR accession number | MW (kDa) | Description | Function |
| --- | --- | --- | --- |
| At5g58230 | 48.2 | MSI1; MULTICOPY SUPPRESSOR OF IRA1 | Chromatin remodelling |
| At3g42170 | 78.8 | DAYSLEEPER | Post-embryonic development |
| At1g06760 | 28.9 | H1.1; histone 1.1 | Nucleosome assembly |
| At3g48750 | 34 | CDC2; CELL DIVISION CONTROL 2 | DNA endoreduplication |
| At1g74560 | 30.6 | NRP1; NAP1-related protein 1 | Nucleosome assembly |
| At1g43190 | 48.2 | PTB3; POLYPYRIMIDINE TRACT-BINDING PROTEIN 3 | Regulation of RNA splicing |
| At3g50670 | 50.4 | U1SNRNP; U1 SMALL NUCLEAR RIBONUCLEOPROTEIN-70K | mRNA splicing |
| At1g80930 | 104 | MIF4G domain-containing protein | mRNA splicing |
| At3g63130 | 58.8 | RANGAP1; RAN GTPASE ACTIVATING PROTEIN 1 | Protein import into nucleus |
| At5g63550 | 59.5 | DEK domain-containing chromatin-associated protein | Chromatin remodelling |
| At3g18790 | 35.4 | Pre-mRNA-splicing factor ISY1-like protein | Generation of catalytic spliceosome for second transesterification step |
| At1g17720 | 56.2 | ATB BETA; protein phosphatase 2A | Cell differentiation |
| At3g29310 | 61.5 | Calmodulin-binding protein-related | Protein folding |
| At5g67630 | 52.1 | ISE4 | Chromatin remodelling |
| At5g55920 | 76.7 | TRM4C, TRNA METHYLTRANSFERASE 4C | Maturation of LSU-rRNA |
| At1g33240 | 74.2 | GTL1; GT-2-LIKE 1 | DNA-templated transcription |
| At4g00238 | 38.2 | ATSTKL1 | DNA-templated transcription |
| At5g41340 | 21.3 | Ubiquitin conjugating enzyme 4 | Protein polyubiquitination |
| At5g18200 | 39 | UTP:galactose-1-phosphate uridylyltransferases;ribose-5-phosphate adenylyltransferases | Galactose catabolic process |
| At5g09850 | 40 | MED26C; MEDIATOR 26C | Transcriptional initiation |
| At2g19570 | 32.6 | CDA1; CYTIDINE DEAMINASE 1 | Cytidine deamination |
| At1g03350 | 51.9 | BSD domain-containing protein | Unknown |
| At2g42810 | 60.2 | PP5; PROTEIN PHOSPHATASE 5 | Chloroplast-nucleus signalling |
| At1g55380 | 75.5 | Cysteine/histidine-rich C1 domain family protein | Unknown |
| At2g22310 | 43.5 | UBP4; UBIQUITIN-SPECIFIC PROTEASE 4 | Protein deubiquination |
| At4g32270 | 27.3 | Ubiquitin-like superfamily protein | mRNA splicing |
| At3g11330 | 55.5 | PIRL9; PLANT INTRACELLULAR RAS GROUP-RELATED LRR 9 | Unknown |
| At2g29570 | 29.2 | PCNA2; PROLIFERATING CELL NUCLEAR ANTIGEN 2 | DNA repair |
| At5g17020 | 123.2 | XPO1A; EXPORTIN 1A | Protein export from nucleus |
| At1g02090 | 29.5 | COP15; CONSTITUTIVE PHOTOMORPHOGENIC 15 | Photomorphogenesis |
| At1g73030 | 22.7 | CHMP1A; CHARGED MULTIVESICULAR BODY PROTEIN/CHROMATIN MODIFYING PROTEIN1A | Embryo development |
| At5g06460 | 119.5 | UBA 2; UBIQUITIN ACTIVATING ENZYME 2 | Protein ubiquitination; DNA damage response |
| At1g20960 | 247 | BRR2A | mRNA processing |
| At1g02080 | 269.7 | NOT1 | Nuclear-transcribed mRNA catabolic process |
| At5g64130 | 15.1 | cAMP-regulated phosphoprotein 19-related protein | Negative regulation of protein dephosphorylation |
| At3g08947 | 96.5 | ARM repeat superfamily protein | Protein import into nucleus |
| At1g78150 | 33.1 | N-lysine methyltransferase | RNA metabolism |
| At5g06460 | 119.5 | UBA2; UBIQUITIN ACTIVATING ENZYME 2 | Protein ubiquitination; DNA damage response |
| At1g63660 | 59.3 | GMP synthase | GMP biosynthetic process |
| At5g06600 | 130.5 | UBP12; UBIQUITIN-SPECIFIC PROTEASE 12 | Protein deubiquitination |
| At3g29075 | 34.4 | Glycine-rich protein | Regulation of gene expression |
| At5g40340 | 114.2 | PDP3, PWWP DOMAIN PROTEIN 3 | Epigenetic regulation of gene expression |
| At4g39200 | 12 | ES25W, RIBOSOMAL PROTEIN ES25W | mRNA binding |
| At1g48410 | 116.4 | AGO1; ARGONAUTE 1 | Regulatory ncRNA-mediated post-transcriptional gene silencing |
| At3g01320 | 156.1 | SNL1; SIN3-LIKE 1 | Negative regulation of transcription |
| At1g67230 | 129 | LINC1; LITTLE NUCLEI 1 | Nucleus organisation |
| At4g38130 | 56 | HDA1; HDA19; HISTONE DEACETYLASE 19 | Histone deacetylation |
| At1g59890 | 132.9 | SNL5; SIN3-LIKE 5 | Negative regulation of transcription |
| At1g70060 | 150.7 | SNL4; SIN3-LIKE 4 | Negative regulation of transcription |
| At5g15020 |  | SLN2; SIN3-LIKE 2 | Negative regulation of transcription |
| At1g24190 | 151.1 | SNL3; SIN3-like 3 | 10 |
| At5g08450 | 104.1 | HDC1, HISTONE DEACETYLATION COMPLEX 1 | Histone deacetylation |

A protein-protein interaction (PPI) network constructed from this subset of proteins shows a suite of well-known partners of HDA19, including five members of the SIN-LIKE (SNL) family. SNL proteins were shown to be involved in the repression of AP2 family transcription factors that repress FLOWERING LOCUS T expression through histone deacetylation (Figure 2C) (Huang et al., 2019). We also identified HDC1, a component of histone deacetylase complexes that interacts with HDA6 and HDA19 (Perella et al., 2016). A bimolecular fluorescence complementation approach revealed that HDC1 binds to the linker histone H1 (Perella et al., 2016), which was identified as a putative interactor of HDA19 in the current dataset. The WD-40 repeat containing protein MULTICOPY SUPRESSOR OF IRA1 (MSI1), a conserved subunit of Polycomb Repressive Complex 2; Xu et al., 2022), and PROLIFERATING CELL NUCLEAR ANTIGEN 2 (PCNA2), involved in DNA replication and damage repair (Xue et al., 2015), were identified as central nodes of the PPI network (Figure 2C).

## Discussion

The current dataset identified a large suite of putative novel interacting partners of a key regulator of plant development and stress responses, HDA19. The identification of SNL members, HDC1, and histone 1 can be considered as validation of the current IP assay. A surprisingly large subset of (predicted) non-nuclear proteins was identified with high confidence, suggesting that some of these proteins may transiently associate with chromatin. Besides expected binding partners such as HCD1, H1, and SNLs, we found that HDA19 interacts with proteins involved in chromatin remodelling, nuclear protein export/import, protein ubiquitination associated with DNA damage repair, and chloroplast-nucleus signalling, suggesting a wide range of largely unexplored functions of HAD19 in etiolated Arabidopsis seedlings.

## Supporting information

Data Set 1

## Dataset description

The mass spectrometry proteomics data have been deposited to the ProteomeXchange Consortium via the PRIDE (Perez-Riverol et al., 2016; Perez-Riverol et al., 2022; Deutsch et al., 2023) partner repository with the dataset identifier PXD045454 and can be accessed through the following link: http://www.ebi.ac.uk/pride/archive/projects/PXD045454.

## Author Contributions

W.S. and I.C.V-B. designed and conceived the experiments. I.C.V-B. conducted the experiments. W.S. and I.C.V-B. analysed the results. I.C.V-B. and W.S. wrote the manuscript.

## Acknowledgments

We thank Chin-Wen Chen and Chuan-Chih Hsu from the IPMB Proteomics Core Laboratory for their help with on-bead trypsin digestion and protein identification. Part A of Figure 1 was created with BioRender.com.

## Supplementary Material

### Data Availability Statement

The datasets generated and analyzed for this study can be found via PRIDE (http://www.ebi.ac.uk/pride/archive/projects/PXD045454).

